# Using a continuum model to decipher the mechanics of embryonic tissue spreading from time-lapse image sequences: An approximate Bayesian computation approach

**DOI:** 10.1101/460774

**Authors:** Tracy L. Stepien, Holley E. Lynch, Shirley X. Yancey, Laura Dempsey, Lance A. Davidson

## Abstract

Advanced imaging techniques generate large datasets capable of describing the structure and kinematics of tissue spreading in embryonic development, wound healing, and the progression of many diseases. These datasets can be integrated with mathematical models to infer biomechanical properties of the system, typically identifying an optimal set of parameters for an individual experiment. However, these methods offer little information on the robustness of the fit and are generally ill-suited for statistical tests of multiple experiments. To overcome this limitation and enable efficient use of large datasets in a rigorous experimental design, we use the approximate Bayesian computation rejection algorithm to construct probability density distributions that estimate model parameters for a defined theoretical model and set of experimental data. Here, we demonstrate this method with a 2D Eulerian continuum mechanical model of spreading embryonic tissue. The model is tightly integrated with quantitative image analysis of different sized embryonic tissue explants spreading on extracellular matrix (ECM) and is regulated by a small set of parameters including forces on the free edge, tissue stiffness, strength of cell-ECM adhesions, and active cell shape changes. We find statistically significant trends in key parameters that vary with initial size of the explant, e.g., for larger explants cell-ECM adhesion forces are weaker and free edge forces are stronger. Furthermore, we demonstrate that estimated parameters for one explant can be used to predict the behavior of other similarly sized explants. These predictive methods can be used to guide further experiments to better understand how collective cell migration is regulated during development.

**Author Summary:** New imaging tools and automated microscopes are able to produce terabytes of detailed images of protein activity and cell movements as tissues change shape and grow. Yet, efforts to infer useful quantitative information from these large datasets have been limited by the inability to integrate image analysis and computational models with rigorous statistical methods. In this paper, we describe a robust methodology for inferring mechanical processes that drive tissue spreading in embryonic development. Tissue spreading is critical during wound healing and the progression of many diseases including cancer. Direct measurement of biomechanical properties during spreading is not possible in many cases, but can be inferred through mathematical and statistical means. This approach identifies model parameters that are able to robustly predict results of new experiments. These methods can be integrated with more general studies of morphogenesis and to guide further experiments to better understand how tissue spreading is regulated during development and potentially control spreading during wound healing and cancer.

## Introduction

Advances in microscopy enable the generation of very large time-lapse datasets over the course of a few hours. Such time-lapse data contains a wealth of information describing both the structures involved in morphogenesis and their kinematics; this information can serve as the input for computational models that allow us to explore the biological and biophysical principles of morphogenesis and to predict the behavior of cells and tissues under a variety of perturbations. This presents many technical challenges, from collecting quality images suitable for machine vision tools to developing computational models that implement appropriate biophysics and physiological rules at useful spatial and temporal scales. To maximize the benefits of these approaches, experimental designs must be able to integrate image data with model simulations in a way that allows robust statistical assessment of model fitness or failure.

Various theoretical and computational frameworks have been used to study tissue spreading and collective cell migration in cases such as wound healing and angiogenesis including reaction-diffusion equations [1–10], continuum mechanical models [11–17], agent-based models [18–22], vertex models [23–27], finite element method (FEM) and CellFIT based inference methods [28,29], and Vicsek and active elastic sheet models [30,31]. These models differ in their assumptions regarding, for example, whether individual cells are distinguishable within the tissue, the basic mechanical nature of the tissue, if cell signaling or mechanical forces are the primary drivers behind cell migration, and whether the models assume a frame of reference that travels with the cells, i.e. a Lagrangian frame of reference, or a fixed frame that cells move across, i.e. an Eulerian frame of reference.

When these frameworks are implemented, they typically estimate physically relevant model parameters through deterministic approaches such as the gradient descent method. Deterministic approaches identify a single set of optimal model parameters that offer a best fit to experimental data [32], however, these approaches do not quantify uncertainty in either the source data or the mathematical model, and thus are ill-suited for comparing parameter fits between two experimental groups. To overcome this limitation, we implement the approximate Bayesian computation (ABC) rejection algorithm [33,34] to estimate biomechanical properties of spreading embryonic tissue via a mathematical model of collective cell migration. While our application is novel in this context, ABC has been used in studying complex biological problems such as those in population genetics, ecology, epidemiology, systems biology, and cell biology [33–36].

While our main goal is to develop a general methodology for combining image data and computational modeling, here we apply this methodology to study epiboly, a form of collective migration, in tissue explants of various initial sizes. Epiboly is a key tissue movement during gastrulation in *Xenopus laevis* embryos, and represents collective cell movements of the ectoderm of the animal cap ectoderm as it spreads and maintains coverage over the exterior of the embryo [37]. We make use of an *in vitro* model of epiboly where ectoderm from the late blastula stage animal cap region of the *Xenopus* embryo is microsurgically dissected and allowed to adhere to a fibronectin-coated substrate (Fig 1A-B). Cultured explants spread over 10 to 24 hours in a manner similar to that observed *in vivo*. Deep cells throughout the ectoderm are thought to contribute to epiboly through adhesion to and traction on a fibronectin coated substrate [38]. Cells at the margin of the explant are thought to generate outwardly directed traction forces that aid spreading. Additionally, as the tissue spreads, the ectoderm decreases in thickness (Fig 1B in [39), but it is not known if this activity is active or merely due to passive strain in the tissue.

**Fig 1.**
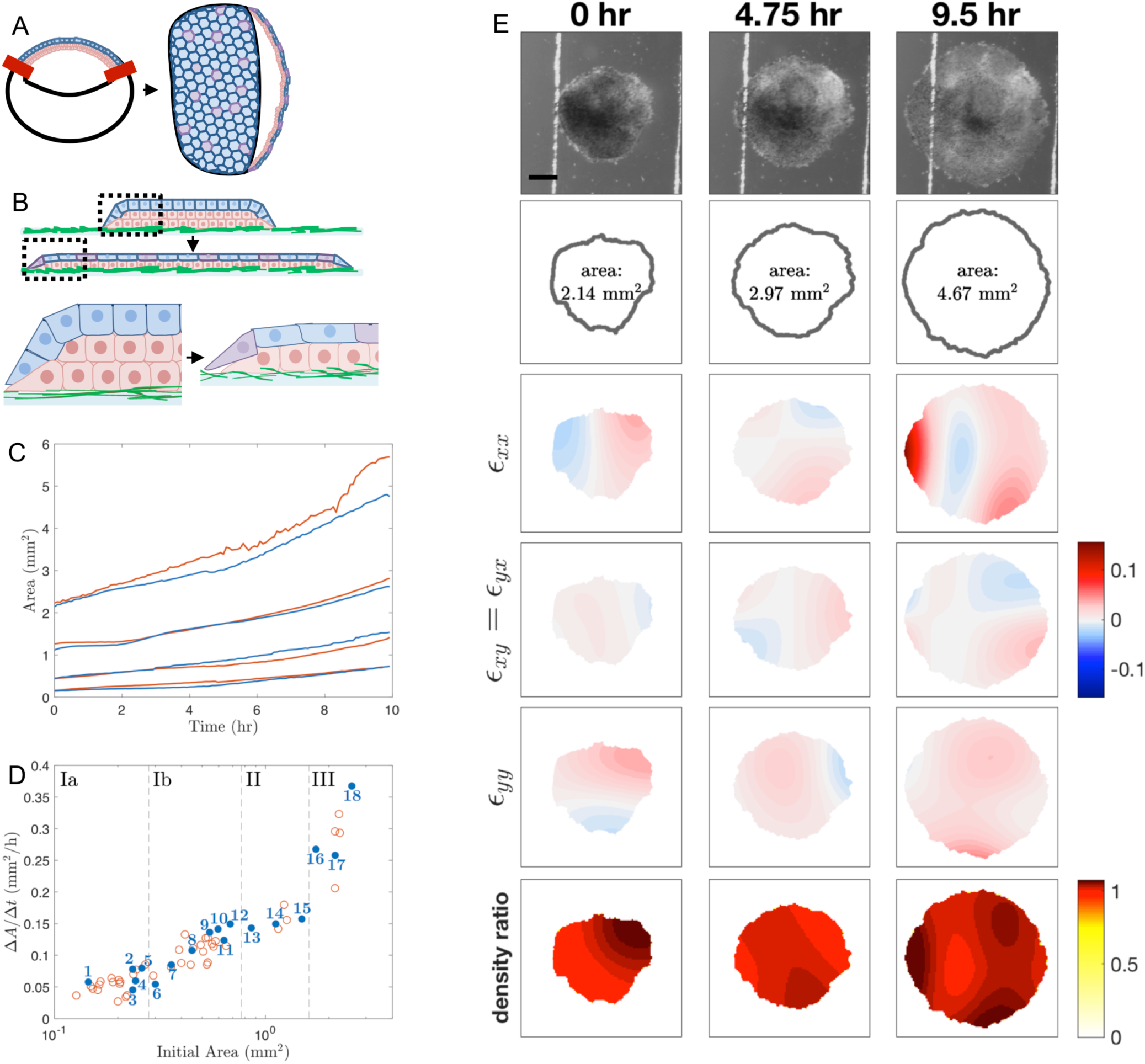
Schematics of experimental setup and image analysis. (A) Schematic of animal cap ectoderm of a *Xenopus laevis* embryo during the late blastula/mid-gastrula stage and at the tailbud stage. At the late blastula stage, a single layer of epithelial cells (blue) lay on top of multiple layers of deep mesenchymal cells (red). The extoderm encloses the blastocoel, a fluid filled cavity in the upper half of the embryo. The red bars indicate where an animal cap explant is cut from the embryo. By the tailbud stage, the ectoderm spreads to cover the entire embryo. The mesenchymal layer undergoes intercalation to become a single layer while the cells in the epithelial layer stretch. (B) Schematic of animal cap ectoderm removed from the embryo during the late blastula through mid-gastrula stages and cultured on fibronectin adsorbed glass (green). As the tissue spreads, deep mesenchymal cells radially intercalate into the epithelial cell layer (purple). Dashed box region shown enlarged below. Note: Experimental images are taken from above, not from the side, so only the pigment-containing apical surface of the epithelial layer is visible. (C) Area of explants over the course of the experiment for representative explants in the model building set (blue) and test set (red). Explants shown here are the same explants analyzed in Fig 4. (D) Average change in area over 10 hours (Δ*A*/Δ*t*), calculated as the slope of the regression line of area measurements over time (as in panel C), for *Xenopus* animal cap explants of different initial sizes. The explants are grouped by initial area into subgroups Ia, Ib, II, and III, indicated by the dashed lines. The blue filled-in circles correspond with the explants included in our model building set and the red circles correspond with the explants included in our test set. The length of time was 10 hours. (E) Time-lapse images of an explant spreading are shown in the first row, followed by the boundary identified by the Level Sets plugin. Strains as defined in Eq. 3 in the S1 File are shown in the third, fourth, and fifth rows. Density ratios (Eq. 15) are shown in the last row. See the S1Video for time-lapse sequences of these still frames. Scale bar: 500 µm.

The physical mechanisms of *Xenopus* embryonic tissue spreading are investigated here by tracking both the movement of the tissue edge and internal tissue deformations. One method of measuring internal deformations of spreading tissue is via cell nuclei tracking [40–42], however, tissues are generally opaque in the *Xenopus* ectoderm, so we adopted an alternative automated method of extracting the local change in texture density between sequential images in time-lapse sequences. This method involves calculating the strain, or local deformation between image pairs using elastic registration [43]. Elastic registration, a form of differential image correlation [44], fits well with an elastic continuum model of tissue spreading, but can also represent deformation in viscous or plastic materials [45].

Our goal in integrating time-lapse image data with a computational model is to determine the extent to which cell shape change, traction forces, adhesion, and tissue elasticity contribute to the mechanics of tissue spreading. In this paper, we adapt an established continuum mechanical mathematical model from Arciero et al. [11] for collective cell migration in which the model parameters correspond to physical quantities. Applying the ABC rejection method to the analysis of epiboly, we identify the distribution of statistically significant parameter sets from multiple experimental data sets of spreading rates and deformation. From these results, we propose tissue mechanical properties such as force production and adhesion are correlated with initial explant size.

## Results

To explore the mechanical processes involved in collective migration during development, we microsurgically isolated ectodermal tissue from the animal cap region of gastrulating *Xenopus* embryos (Fig 1A). During gastrulation, this tissue spreads to cover the exterior of the embryo. Once isolated and placed on a fibronectin-coated substrate, tissue explants spread radially outward [46,47] (Fig 1B). The initial area of the explant affects the spreading rate with larger explants spreading faster than smaller ones (Fig 1C-D), suggesting that the initial area of an explant affects the mechanics of the tissue. To understand the source of these mechanical interactions, we sought to integrate a mathematical model of epiboly with kinematic data on tissue boundaries and internal strains from time-lapse data (Fig 1E; Materials and Methods). Through parameter estimation and statistical techniques, we correlated the effect of an explant’s initial area on estimated parameter values in the model.

Our experimental data set contained 59 explants from two clutches of eggs; these explants were shaped so the initial areas varied eighteen-fold (Fig 1D). We grouped these explants into four subgroups based on similar initial areas, where groups Ia and Ib are comprised of the smallest explants and the largest explants constitute group III (Fig 1D). The cutoffs between groups were chosen in order to distribute explants evenly where large natural breaks in measured area occurred. By grouping, we hoped to expose variation between individual explants as well as trends over the size-grouped explants. We chose eighteen representative explants for our model building and left the remaining explants to test the predictive power of the method. Note: explants were named in numerical rank by their initial area (see Fig 1D and S1 Table; e.g. the smallest explant is named Explant #1).

## Mathematical Model

To study explant spreading, we modify the two-dimensional (2D) cell migration model of Arciero et al. [11], which is derived from mechanics principles based on adhesion between the tissue and substrate, elastic coupling of cells within the tissue, and forces generated by lamellipodia. This model also recapitulates *Xenopus* explant mechanics and movements captured by image analysis.

The model of Arciero et al. [11] was originally applied to small intestinal epithelial (IEC-6 cell) and Madin-Darby canine kidney (MDCK) epithelial cell layers. Minor modifications to the model are required to apply to explant spreading since the biological mechanisms of material growth in the visible cell layer are different. For example, epithelial cell layers in MDCK and IEC-6 sheets are one cell thick and exhibit volumetric growth through cell proliferation. By contrast, while cells in *Xenopus* embryos divide, the whole embryo does not change total volume from fertilization to feeding tadpole stages [48]. Furthermore, the animal cap is a composite tissue consisting of a single epithelial layer underlain by one or more layers of deep mesenchymal cells. Spreading in the animal cap can undergo apparent 2D volumetric growth but this movement is contiguous with phases in which cells move from one layer to another and change in thickness. Since the time-lapse images of explant migration are taken normal to their plane of movement, we model the explant tissue as a single layer that can be tracked using only the cells in the visible epithelial layer of the tissue. Like models of MDCK and IEC-6 sheets. we also assume that the explant is uniformly thick, but note that changing this condition would be a useful future extension.

In the model of Arciero et al. [11], a 2D single cell layer is represented by an elastic continuum capable of deformation, motion, and cell proliferation. Since the multicellular ectoderm of *Xenopus laevis* animal caps is continuous and does not fracture or shear, a continuum model of tissue migration is appropriate [49]. The tissue is represented as a 2D compressible inviscid fluid and the motion of cells is described in spatial (Eulerian) coordinates. The parameters in this model correspond to physical properties in the tissue, and thus we can deduce the material properties of the tissue that best represent observations from a single experiment. The governing equation of the model of Arciero et al. [11] is

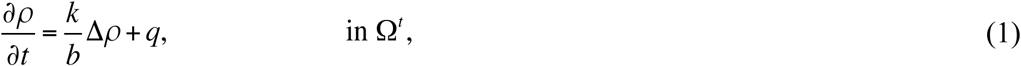

where the variable *ρ*(**x**,*t*) describes the tissue density as a function of position **x**=(*x,y*) and time *t, k* and *b* are parameters described in Table 1, *q* is a growth term, and Ω^*t*^ is the domain that describes the extent of the tissue at time *t* (Fig 2A-B).

**Table 1.**
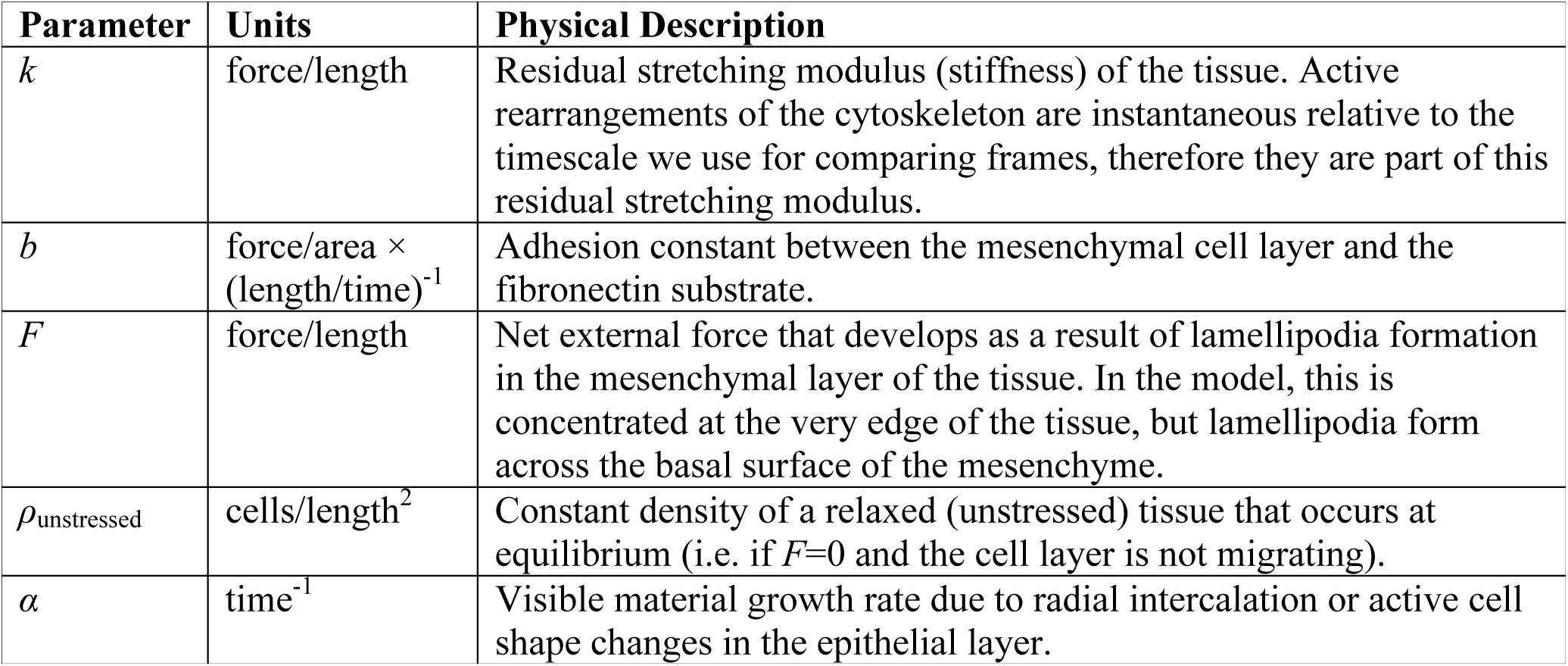
Mathematical model parameters, units, and descriptions.

**Fig 2.**
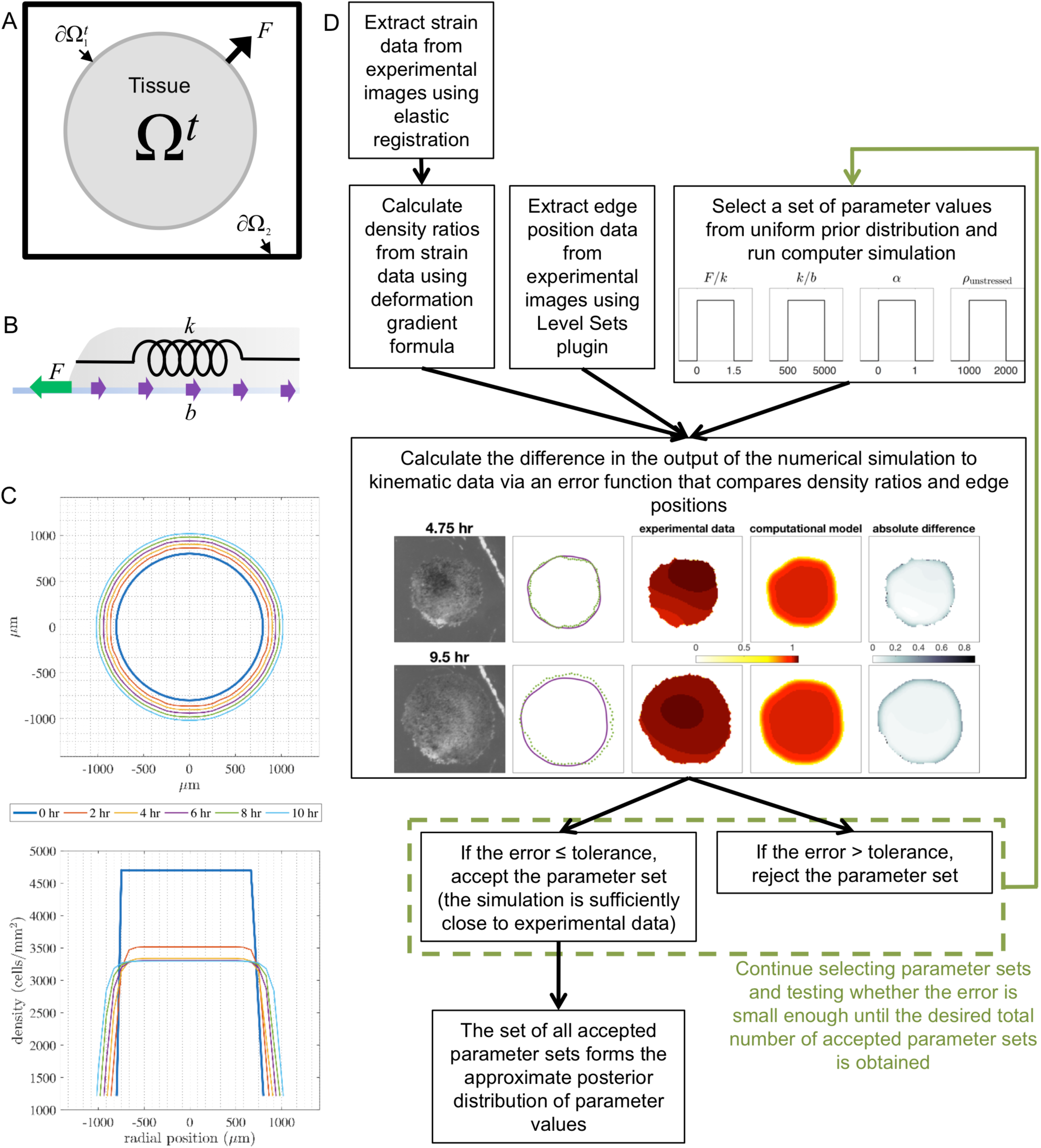
Schematics of the numerical simulations and approximate Bayesian computation rejection method. (A) Computational domain of the moving boundary initial value problem as if looking at the animal cap explant from above. The tissue is located in the shaded region (Ω^t^), the tissue boundary is denoted by ∂Ω_2_^*t*^, the edge of the computational domain is denoted by ∂Ω_2_, and *F* is the net external force per unit length at the tissue boundary that develops as a result of lamellipodia formation. (B) Schematic of a cross-section of the explant during spreading and the forces governing its movement. The green arrow represents the lamellipodia force *F*, the purple arrows represent the adhesion force *b*, and the spring represents the residual stretching modulus of the tissue *k*. (C) Toy simulation showing the output of the boundary locations (as viewed from above) and density profiles (as a cross-section through the horizontal axis) every 2 hours over 10 hours. The fixed grid used is shown as gray dashed lines. (D) Flow chart of the approximate Bayesian computation (ABC) rejection algorithm implemented for parameter estimation.

The boundary conditions and initial condition are given by

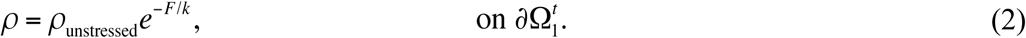

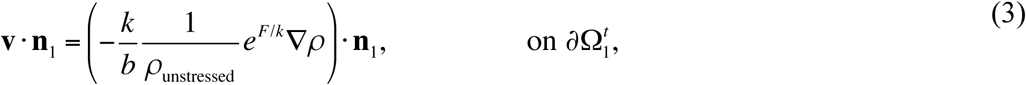

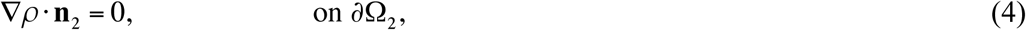

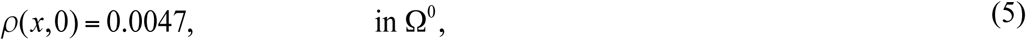

where *ρ*_unstressed_ and *F* are parameters described in Table 1, **v**(**x**,*t*) is the velocity of the tissue, **n**_1_(**x**,*t*) is the outward unit normal to the tissue boundary ∂Ω_1_^*t*^, and **n**_**2**_(**x**,*t*) is the outward unit normal to the edge of the computational domain ∂Ω_2_. We note that in our simulations, the computational domain ∂Ω_2_ is large enough so that the tissue boundary ∂Ω_1_^*t*^ is enclosed by ∂Ω_2_ . See the S2 File for further details on the derivation of Eq. 1 and the boundary and initial conditions.

In the original model of Arciero et al. [11], the growth term *q* was included in Eq. 1 to model the volumetric proliferative growth which is typical for spreading sheets of cultured cells, such as IEC-6 or MDCK cells, and taken to be logistic growth. As discussed above, total embryonic volume is conserved even as cells divide. Therefore, each explant initially contains a fixed number of cells that can divide but their proliferation does not increase the net volume of the explant (a process more generally termed reductive cleavage). Thus, spreading of explants does not depend on volumetric growth provided by cell proliferation [48].

While Xenopus explants do not undergo volumetric growth, their surface area can be increased by cells moving from one layer to the other or by cell height changes. The process of radial intercalation in *Xenopus* ectoderm allows cells to move from the deepest layers into the more superficial layers, a process that plays no role in changing the surface area of single-layered cultured cell sheets. Empirically, we estimate that there is at most a 50% more cells on the surface of the initial explant from either active shape changes or intercalated cells from the lower mesenchymal layer, which is a reasonable upper bound based on the estimate that no more than 20% of the surface cells in the embryo come from the deep mesenchymal layer [38]. Therefore, we modify *q* in Eq. 1 to represent surface material added through other means, such as active cell shape change or radial intercalation. After qualitatively testing growth functions, we chose a mass-limited density-dependent logistic growth function to describe the addition of visible material into the surface layer,

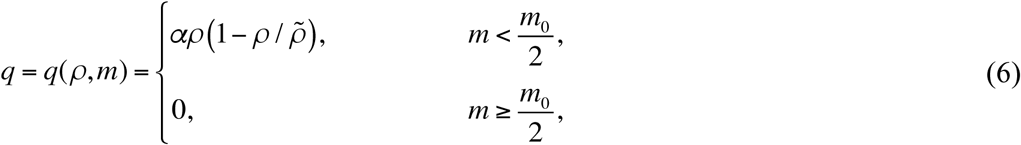

where *α* is a parameter described in Table 1, 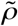 is the limiting density assumed to correspond to a compressed tissue such that 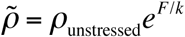 [11], *m(t)* is the cumulative amount of mass that has been added to the visible surface layer through time *t*, and *m*_0_ is the initial mass.

Details of the numerical method implemented to solve the model equations may be found in Arciero et al. [11]. A simulation of a simple case of tissue spreading illustrates how the tissue boundary and internal tissue density change over time (Fig 2C).

### Parameter Estimation Using Approximate Bayesian Computation (ABC)

To infer material properties of the tissue that are not directly measurable and predict tissue behavior we use the approximate Bayesian computation (ABC) rejection method [33,34]. ABC determines probability distributions for the model parameters for each individual tissue explant (see Fig 2D for a flow chart depicting this process). The results from individual explants from our model building set are then compared to determine how parameter values for explants vary across a range of explant sizes (Fig 1D).

Estimating parameters for our model in a deterministic manner appeared infeasible as preliminary studies suggested that our model had parameter structural non-identifiability, or functionally related model parameters [50]. While parameter identifiability has been explored extensively for ordinary differential equations, for example determining whether different parameter sets give rise to different model predictions, it has not been studied as deeply in partial differential equation models [51]. Thus, from preliminary analysis of parameter space and the knowledge that models similar to our own have shown non-identifiability relations between parameter values [52], we adopted ABC rejection because it allows for quantification of the uncertainty of our parameter estimates. Furthermore, quantifying uncertainty in single data sets permits comparison between estimated parameter values among multiple data sets.

To use ABC rejection, the first step is to specify a prior distribution for each of the parameters. This prior distribution is our best educated guess at how parameter values *F, k, b, α*, and *ρ*_unstressed_ (Table 1) vary. However, since *F, k*, and *b*, appear in the model equations only as the ratios *F/k* and *k/b*, we cannot uniquely determine each parameter and thus only consider them as ratios. The prior distribution for each parameter is a broad uniform distribution *U*(*a,b*) with probability density function defined by *f*(*x*) = 1/(*b*-*a*) for *x* ∈ [*a, b*], and 0 otherwise, where *F*/*k* ∼ *U*(0,1.5) [dimensionless], *k*/*b* ∼ *U*(500,5000) [µm^2^/h], *α* ∼ *U*(0,1) [h^-1^], and *ρ*_unstressed_ ∼*U*(1000,2000) [cells/µm^2^]. We chose the prior distribution from previous parameter estimates for IEC-6 and MDCK cell sheets [11,14] and our estimates of Xenopus ectoderm tissue density, which is determined from observed proliferative behavior and average cell size.

The second step of ABC rejection is to sample randomly from the distribution of each of the parameters. From each sampling we obtain a single set of parameters {*F*/*k, k*/*b, α, ρ*_unstressed_}, which we use in a simulation of the mathematical model. Quantification of how well the numerical simulation corroborates with experimental data is determined by calculating the ABC Error Function (see Materials and Methods) from the sum of the mean-squared difference between the observed and simulated density ratios over the time-course of the simulation.

We collected 10,000 parameter sets that result in total error (Eq. 12) less than or equal to a tolerance threshold of 1500 for each explant. The computational expense of running this parameter set collection on a supercomputer was over 100,000 CPU hours limiting the feasibility of collecting thousands or millions of additional samples. We retained all of the 10,000 parameter sets to be posterior samples to represent the distribution of parameter values that are optimal, however, some analyses (indicated below) were carried out with the smallest 20% (in terms of the total error) of the 10,000 parameter sets.

For each of the eighteen explants, we used the ABC rejection algorithm to obtain posterior parameter value distributions for *F*/*k, k*/*b, α*, and *ρ*_unstressed_. To visualize the posterior distributions for the four parameters, we projected the distributions down to one and two dimensions (see Fig 3A for one explant). Along the diagonal are smoothed histograms for the one-dimensional projections of the individual parameters. Below the diagonal are the two-dimensional projections of pairs of parameters. Darker shaded areas correspond with higher frequency where higher frequency corresponds with the parameter value being sampled in the posterior distribution more often.

**Fig 3.**
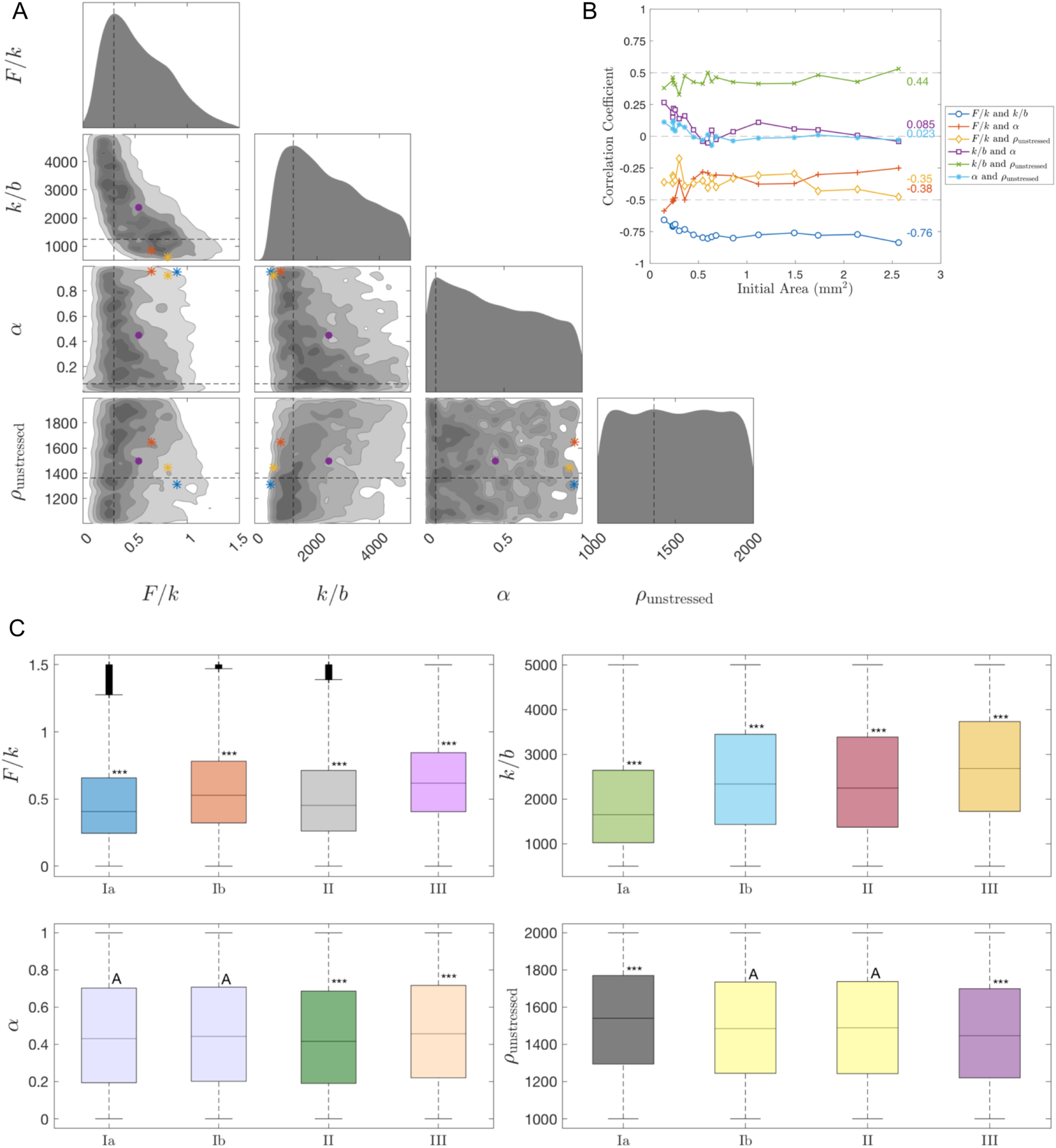
Results of approximate Bayesian computation rejection method and statistics results. (A) Triangle plot of posterior distributions obtained by the ABC rejection method for Explant #8. Along the diagonal, the plot shows smoothed histograms for the 1-D projections of the parameters *F*/*k, k*/*b, α*, and *ρ*_unstressed_. The lower part of the triangle shows the 2-D histograms for all pairs of these four parameters. The darker shades in the 2-D histograms denote more frequently occurring values of parameters, and the dashed vertical and horizontal lines denote the most frequently occurring values of each parameter. The purple circle indicates the mean parameter sets and the blue, red, and yellow asterisks indicate the parameter sets with the smallest errors, respectively. 10,000 parameter sets were used. Cf. S1 Fig for the triangle plot colored in terms of the error (Eq. 12) instead of frequency. (B) Correlation coefficient for each pair of parameters for all 18 explants. The average correlation coefficient for each pair of parameters over all the explants is displayed on the right side of the plot. The smallest 20% (in terms of the total error) of the 10,000 parameter sets were used per explant. (C) Boxplots of the 1-D projections of the posterior distributions of the parameters *F*/*k, k*/*b, α*, and *ρ*_unstressed_ for all 18 explants, grouped by region Ia, Ib, II, and III. The compact letter display (CLD) and colors reflect the results for Tukey’s multiple comparisons test; in particular, boxplots labeled with the same letter were considered not statistically different (based on significance level 0.05). Boxplots labeled with *** were statistically significantly different than all the other boxplots. Coloring was chosen to reflect groups of boxplots that were labeled similarly. The plus signs indicate outliers, the lines within the boxes indicate the median, and the length of the boxplot indicates the interquartile range. 10,000 parameter sets were used per explant. Cf. S2 Fig for the non-grouped boxplots.

Analysis of parameters describing explant spreading reveals an inverse relationship between *F/k* and *k/b*, which implies non-identifiability, and less distinct relationships between the other pairs of parameters, which may imply identifiability or weak non-identifiability (Fig 3A, see also S1 Fig). To verify the association between the different parameters, we calculated the correlation coefficient for each pair of parameters for each explant (Fig 3B) using the smallest 20% (in terms of the total error) of the 10,000 parameter sets per explant. A correlation coefficient of *r*=+1 or *r*=-1 corresponds to two parameters being strongly correlated, a correlation coefficient between 0.5<|*r*|<1 is moderately to strongly correlated, and a correlation coefficient between 0<|*r*|<0.5 is weakly to moderately correlated. We found that *F/k* and *k/b* were strongly negatively correlated, pair *k/b* and *α* and pair *α* and *ρ*_unstressed_ were weakly correlated, and that the other parameter pairs were moderately correlated.

We also calculated 95% confidence intervals for the one-dimensional projections of the posterior distributions for each parameter for each explant (see S1 Table). We observe apparent trends in the mean parameter value versus the initial area of the explant, but to verify this we ran ANOVA tests. Using 10,000 parameter sets per explant and grouping them by initial size into groups Ia, Ib, II, and III, we determined that the p-value was strictly less than 10^−29^ for all the parameters *F*/*k, k*/*b, α*, and *ρ*_unstressed_. This implies that mechanical parameter values differed for the four groups of explants. We repeated this analysis without grouping the explants by size and found even smaller *p*-values.

Additionally, we ran Tukey’s multiple comparisons test to compare the average parameter values of the eighteen explants, grouped by initial area and individually, and to determine which showed statistically significant differences. Fig 3C shows boxplots of the one-dimensional projections of the posterior distribution for each parameter by initial size. The boxplots are labeled with a compact letter display (CLD) such that boxplots labeled with the same letter were not statistically different (based on significance level 0.05). In S2 Fig, boxplots of the non-grouped explants are shown. We found that *α* and *ρ*_unstressed_ are not strongly affected by initial area (denoted by more overlapping letters in the CLD as well as boxplots of the same color), but *F/k* and *k/b* are affected.

To further test whether estimated values of *F/k, k/b, α*, and *ρ*_unstressed_ are associated with the initial area, we ran a linear regression *t*-test with the mean values of each parameter from the one-dimensional projection of the posterior distribution for each explant. We calculated that *p*-value *p* = 0.02 for parameter *F/k, p* = 0.002 for *k/b*, and *p* = 0.0004 for *ρ*_unstressed_, which suggests a relationship between the parameter value and the initial explant area, yet, we found that *p* = 0.17 for *α*, which reduces the likelihood of such a relationship. Thus, the material growth rate of the spreading tissue, *α*, is the least variable parameter among explants of various sizes.

The slopes of the sample regression lines for the mean value of *F/k* vs. initial area and the mean value of *k/b* vs. initial area are both positive while the slope for the mean value of *ρ*_unstressed_ vs. initial area is negative. This suggests a general trend that both *F/k* and *k/b* increase as the initial area increases and *ρ*_unstressed_ decreases as the initial area increases. Furthermore, if we assume that the stretching modulus *k* is constant for each explant, which is a realistic expectation since explants are isolated from similar locations, then these results imply that lamellipodia forces (*F*) are stronger and the adhesion forces (*b*) are weaker in large explants. A similar relationship between the lamellipodia force and the size of the initial explant was also observed in primary mouse keratinocyte cell colonies [53,54], where traction stresses increase with colony radius. In addition, from the inverse relationship between *F/k* and *k/b* in the triangle plot (Fig 3A), we predict that if adhesion is increased, then the measured traction forces will also increase.

### Prediction and Method Robustness

To determine whether the posterior distributions for *F/k, k/b, α*, and *ρ*_unstressed_ were robust, we took the parameter set that resulted in the smallest error (Eq. 12) for an explant in our model building set and used it to simulate an explant in the test set with a similar initial area (Fig 4A-D) or was from a different size group (Fig 4E-H). For similarly sized explants we found the error (Eq. 12) remained small, but for differently sized explants the error was large. Furthermore, we observed that similarly sized explants that do not have similar average changes in area over time (Δ*A*/Δ*t*), such as the areas of Explant #8 and Explant B which are represented in the third and fourth lowest curves in Fig 1C, also resulted in larger errors. Hence, our posterior distribution results are robust since explants of similar size will have similar model parameters that best fit the experimental data whereas explants of different size will not yield a good fit. (Note: for comparison, simulations for Explants #1, 8, 14, and 17 are shown in S3 Fig and the S9-S12 Videos.)

**Fig 4.**
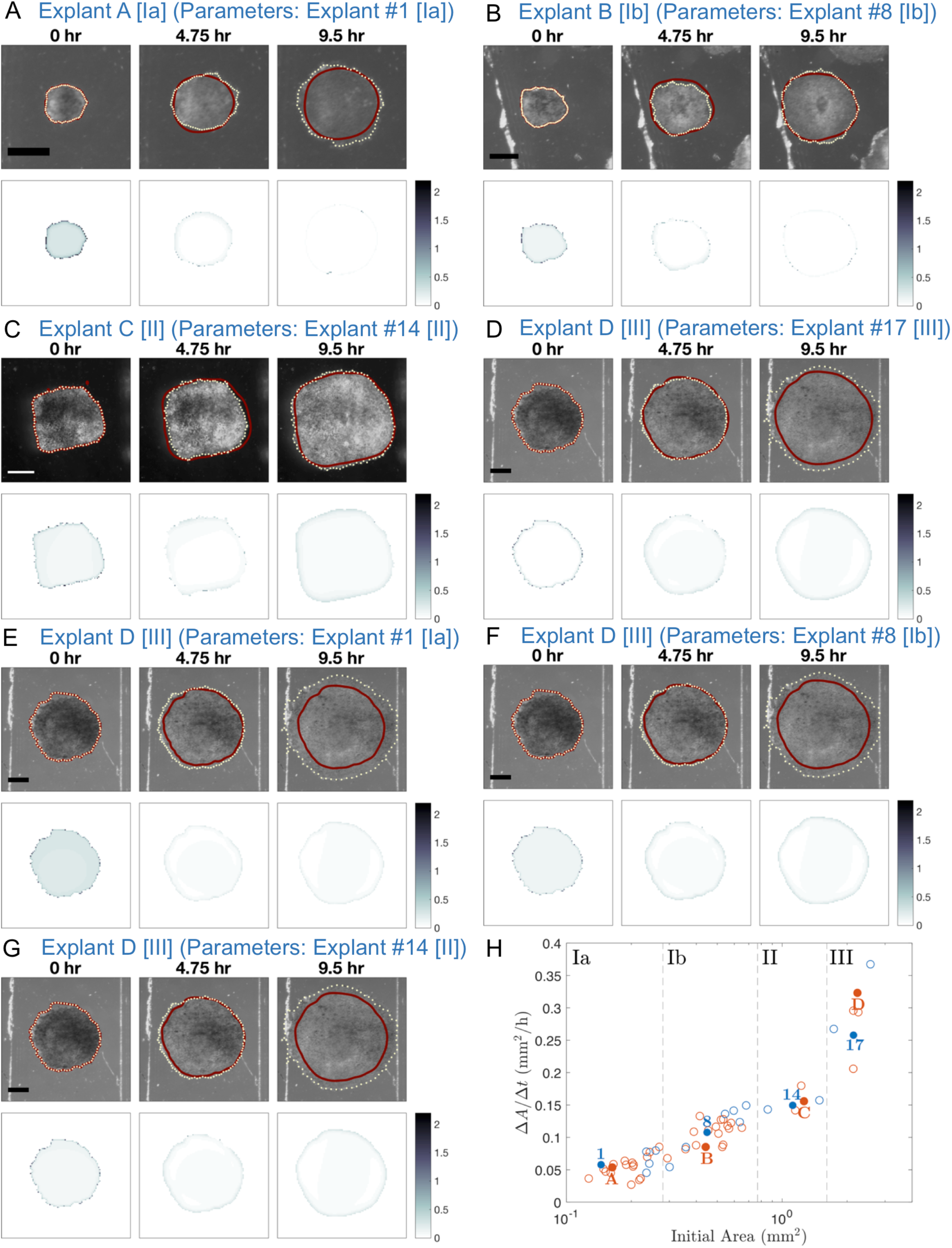
Simulation of tissue spraeding for various test explants using estimated parameter values from similarly sized or differently sized explants. Progression of *Xenopus* animal cap explant tissue spraeding at 4.75 hour time intervals for test explants A, B, C, and D with the parameter set that resulted in the smallest error (Eq. 12) from the posterior distributions for a similarly sized explant from the same region (A-D) or a differently sized explant from a different region (E-G). In the top panel, the computed edge from the mathematical model is represented by a solid dark red curve and the experimental edge is represented by a dotted light yellow curve. The bottom panel shows the absolute value of the density ratio at each grid node (where both density ratios are nonzero) of the experimental data between the given time point and 25 minutes later minus the computed density ratio. Scale bar: 500 µm for each explant. See the S2-S8 Videos for time-lapse sequences of these still images. (A) Test Explant A [Region Ia] (initial area 0.16 mm^2^) using parameters for Explant #1 [Region Ia] (initial area 0.14 mm^2^): *F/k* = 0.6101, *k/b* = 510 µm^2^/h, *α* = 0.9479 h^-1^, *ρ*_unstressed_ = 1404 cells/µm^2^. The total error with these parameters for Test Explant A is 1091 while for Explant #1 it is 1122. (B) Test Explant B [Region Ib] (initial area 0.44 mm^2^) using parameters for Explant #8 [Region Ib] (initial area 0.45 mm^2^): *F/k* = 0.9059, *k/b* = 510 µm^2^/h, *α* = 0.9492 h^-1^, *ρ*_unstressed_ = 1313 cells/µm^2^. The total error with these parameters for Test Explant B is 1109 while for Explant #8 it is 820. (C) Test Explant C [Region II] (initial area 1.26 mm^2^) using parameters for Explant #14 [Region II] (initial area 1.12 mm^2^): *F/k* = 0.7948, *k/b* = 635 µm^2^/h, *α* = 0.9410 h^-1^, *ρ*_unstressed_ = 1530 cells/µm^2^. The total error with these parameters for Test Explant C is 869 while for Explant #14 it is 774. (D) Test Explant D [Region III] (initial area 2.23 mm^2^) using parameters for Explant #17 [Region III] (initial area 2.14 mm^2^): *F/k* = 0.8295, *k/b* = 950 µm^2^/h, *α* = 0.8963h^-1^, *ρ*_unstressed_ = 1752 cells/µm^2^. The total error with these parameters for Test Explant D is 933 while for Explant #17 it is 834. (E) Test Explant D [Region III] (initial area 2.23 mm^2^) using parameters for Explant #1 [Region Ia] (initial area 0.14 mm^2^): The total error for Test Explant D is 1486. (F) Test Explant D [Region III] (initial area 2.23 mm^2^) using parameters for Explant #8 [Region Ib] (initial area 0.45 mm^2^): The total error for Test Explant D is 1136. (G) Test Explant D [Region III] (initial area 2.23 mm^2^) using parameters for Explant #14 [Region II] (initial area 1.12 mm^2^): The total error for Test Explant D is 1140. (H) Average change in area over time (Δ*A*/Δ*t*) for Xenopus animal cap explants of different initial sizes (cf. Fig 1D). The blue numbers represent explants from our model building set and the red letters correspond with the explants selected from our test set

## Discussion

Time-lapse imaging of tissue explant spreading generates complicated data sets that are information-rich. Here, we present a formal methodology for extracting mechanical mechanisms of tissue spreading in differently sized embryonic tissue explants by integrating complex image information with a mathematical model of cell migration. In order to directly relate the experimental kinematics of spreading of multiple tissue explants to mechanical properties of the tissue, we focused on one mathematical model, the previously-developed Eulerian mechanical model of Arciero et al. [11], that represents forces involved in migration within a 2D elastic continuum. In this modeling framework, extended here to represent *Xenopus* animal cap epiboly, we can directly couple experimentally derived kinematic data to model parameters that correspond to physical properties in the tissue.

Taking advantage of naturally abundant pigment patterns allowed us to map strain across the tissue and use changes in area strain to estimate local changes in tissue density. While we used variations in pigment pattern to calculate strain and time-dependent changes in density, any landmark-rich image sequence of sub-cellular organelles, dye, or fluorescence could be used to obtain similar density ratios. Furthermore, our assumption that the tissues are elastic for the image registration and the mathematical model could be relaxed to accommodate more realistic constitutive models such as viscoelasticity or viscoplasticity [55–57]. Since strain mapping techniques do not depend on free edges, our method can also be extended beyond tissues in two dimensions to analyze more complex structures, organs, and whole organisms.

Previous studies of tissue spreading by Arciero et al. [11] and Mi et al. [14] used two different cultured cell lines, rat intestinal epithelial cells (IEC-6) and Madin-Darby canine kidney (MDCK) epithelial cells, and estimated model parameters for individual experiments. While these studies report one parameter set that matches a single set of experimental data, we show here that while there may be a single “optimal” parameter set for each set of experimental data, due to the inverse relation between model parameter ratios *F/k* and *k/b*, there is a wide range of “close to optimal” parameter sets. Thus, to compare different sets of experimental data, we needed to implement a robust statistical analysis capable of accounting for the uncertainty in the model parameter estimates.

To estimate this uncertainty we used the approximate Bayesian computation (ABC) rejection method to collect multiple parameter sets that resulted in sufficiently good fits to the experimental data. The goodness of fit was determined via an error function that calculated the difference between the model-generated and image-based measurements of tissue boundary and density changes throughout the simulation, rather than just matching a single end time point. Hence, we used information from entire time-lapse sequences to estimate physically-relevant parameter values. Statistical analysis then allowed us to examine the difference in means in multiple parameter set distributions for explants of varying sizes. Using this methodology, we found interdependencies between parameters *F*/*k* and *k*/*b*. Such interdependencies may reflect physiological changes in the tissue, for instance, the cells’ ability to modulate traction or adhesion strength to maintain spreading behavior in the face of perturbations in their mechanical properties or microenvironment. If we assume that the stretching modulus *k* is fixed among explants, we find trends suggesting lower adhesion and larger traction forces are present in larger explants. For reasons that are not clear to us, smaller explants required more simulation runs to identify best fit states than larger explants (S4 Fig) and may reflect a more limited range of good-fit parameters. Interestingly, the material growth rate, *α*, is the least variable of all the parameters and may reflect an invariant developmental program for cell intercalation and cell shape change across a range of explant sizes. Thus, from the results of our analysis, the driving forces of tissue are strong net external forces that develop as a result of lamellipodia formation and weak adhesion (traction) forces between the tissue and the substrate.

High computational expense limited the number of parameter sets gathered for each explant to 10,000 for the ABC method as well as the size of our model building set to 18 of 59 available explants. However, we observed that the parameter values found were robust among similarly sized explants (Fig 4). Indeed, for explants with similar average change in area over time (Δ*A*/Δ*t*), simulations with a parameter set that resulted in a small error for a similarly sized explant in the test set. Thus we would expect that the trends we discovered would be the same if we analyzed more explants in the future, taking into account embryo to embryo variation in stiffness [58], clutch to clutch variation [59], and possible environmental variations such as room temperature that might affect rates and deformation maps of tissue spreading [60].

The probability distributions provide ranges for parameters *F*/*k* and *k*/*b* that can be compared to direct measurements (Fig 4). Since parameters *F, k*, and *b* appear only as the ratios *F*/*k* and *k*/*b* in the mathematical model, to estimate values for each of those parameters individually we would need a measurement of at least one of the quantities. From previous studies we estimate that the stiffness of the ectodermal epithelial layer of *Xenopus* gastrula is approximately 1.2 mN/m [55,59,61]. Using this value as an approximation for *k*, keeping in mind that our explants included both the epithelial and mesenchymal layers, we predict that traction forces at the explant edge (*F*) range from 0.59 to 0.91 nN/µm and that adhesion (*b*) ranges from 0.0004 to 0.0007 nNh/µm^3^ (using the minimum lower bound and maximum upper bound of the 95% confidence intervals in S1 Table).

Our work suggests future extensions to the Eulerian model include incorporating spatial and temporal heterogeneity within the tissue and in its environment in order to model geometrically and mechanically complex tissues in diverse microenvironments that change over time. For example, complex features that might improve agreement with asymmetrically spreading tissue include allowing parameters to change over time, tissue viscosity, anisotropic tissue stiffness, patterned extracellular matrix, and distinct material models for substrate non-linear elasticity. Furthermore, since the *Xenopus* animal cap explants consist of a single epithelial layer and multiple layers of deep mesenchymal cells, models that explicitly represent interactions between cell layers might better account for their integrated spreading movements.

Recent years have witnessed a deluge of high quality image data from confocal and light-sheet microscopy from both *in vivo* and *ex vivo* models of tissue morphogenesis, but efforts to integrate such kinematic descriptions with mechanical models have lagged behind systematic methods that are capable of inferring patterns of force production and patterns of tissue mechanical properties. Our Eulerian mechanical model with approximate Bayesian computation achieves that goal, suggesting future mechanical modeling approaches could be directly integrated with kinematic data from multiple experiments. These efforts will lead to more rapid analysis of morphogenetic movements and the improved correlations between molecular scale perturbations and phenotypic change.

## Materials and Methods

### Embryos and sample preparation

Eggs were collected from *Xenopus laevis* frogs and fertilized *in vitro* using standard methods [48]. After fertilization, eggs were dejellied and cultured in 1/3 X Modified Barth’s Saline (MBS) [62]. Embryos used to analyze intercalation were transferred to a 3% ficoll solution in 1X MBS and were injected (Harvard Apparatus) with approximately 1ng mRNA encoding a membrane tagged GFP (mem-GFP [63]) at the two cell stage, after which they were returned to 1/3 X MBS. Embryos were raised to approximately Stage 10 [37] and were transferred to explant culture medium (Danalchik’s for Amy; DFA) [64]. Embryos were devitelinized with forceps and the animal cap ectoderm was microsurgically removed using hair tools (Fig 1A-B). Tissue explants for brightfield were cultured on a sterile petri dish (Fisher) or for confocal in a alkaline-washed glass-bottomed chamber coated with human plasma fibronectin (25 µg/mL at 4°C overnight in 1/3 X MBS; Roche Molecular Biochemicals) in DFA supplemented with antibiotic/antimycotic (Sigma). The fibronectin coating concentration was chosen because it shows similar spreading results over a broad range of concentrations (S5 Fig). Fibronectin-coating was assumed to be homogeneous across the substrate. Explants were allowed to adhere at least 30 minutes prior to imaging.

After an initial adhering stage explants generally spread radially outward maintaining roughly their initial shape. We note that fibronectin coating may be disrupted or inadvertently scratched when the tissue explants are physically manipulated and positioned for culture. We suspect such defects were present in the infrequent cases where spreading was inhibited in a region or strongly asymmetric and have excluded those cases. We cannot formally rule out fibronectin concentration-dependent effects on migration rates, however, explants spread at similar rates over a broad range of fibronectin coating concentrations and are only significantly impacted when coating is carried out below 1.0 µg/mL (S5 Fig).

Animals used in this study were treated according to an animal use protocol issued to Dr. Davidson (IACUC Protocol #12020250). This protocol covers production and the specific use of embryos and organotypic explants described in this study and has been reviewed and approved by the University of Pittsburgh Institutional Animal Care and Use Committee (Assurance #: A3187-01) in order to meet all US government requirements.

### Imaging

Brightfield stereoscope images of tissue explants spreading were collected every 5 minutes using a CCD camera (Scion Corporation) mounted on a variable zoom stereoscope (Stemi 2000; Zeiss) (Fig 1E; first row). Explants were imaged with side- or grazing-illumination with a fiber optic lamp (Fostec) against either a grey or black background. Multiple explants could be imaged simultaneously using a motorized XY stage (Marzhauser and Ludl) controlled through microscope automation and image acquisition software (µ-Manager [65]). Images of a 1 mm stage micrometer were collected after each experiment to calibrate spatial dimensions in the time-lapse images. All explants were recorded for at least 10 hours before overt differentiation occurs. We chose a 10-hour time frame because the cells within tissue explants will differentiate until yolk nutritive stores run out (approximately 3 days after explanting). From longer time-lapse videos, we typically observe that spreading continues for 20 to 24 hours beyond our analysis, even as ectoderm cells differentiate as larval epidermis.

In order to determine absolute cell density, high resolution confocal images of membrane tagged GFP in live samples were collected using a 20x dry objective (Leica HCX PL APO; n.a. 0.70) on an inverted compound microscope (DMI6000) equipped with a laser-scanning confocal scanhead (SP5; Leica Microsystems). Confocal images were collected with the pinhole set to 1 airy unit. To evaluate the apical epithelial cell density, we collected either a single image or a short stack of images over a z-range of 50 to 100 µm. In the case where z-stacks were collected, a single image frame was prepared for segmentation after a maximum intensity projection.

### Image Analysis

Subsequent image analysis of time-lapse sequences were performed using ImageJ ([66]; available for download at http://imagej.nih.gov/ij/) and Fiji ([67]; available for download at http://fiji.sc/) as described below. Cell boundaries were identified semi-automatically using SeedWater Segmenter ([68]; available for download at https://my.vanderbilt.edu/shanehutson/software/seedwater-segmenter/) in order to estimate cell density.

Tissue edge boundaries were identified from single stereoscope images using the Level Sets plugin in Fiji [67] (Fig 1E; second row). The error is expected to be less than 5 pixels, which corresponded to no more than 2 cell widths (approximately 40 µm). To calculate the average area spreading rate, the area within this boundary was determined at each time point. These areas were then fit using linear regression. The slope of the resulting regression line was defined as the average change in area over time (Δ*A*/Δ*t*) (Fig 1D).

### Approximate Bayesian Computation (ABC) Error Function

To identify suitable simulation parameter sets, we constructed an error function from the sum of the mean-squared difference between the observed and simulated tissue edge positions and interior strains. For the error in edge positions, we calculated the minimum distance from each computational edge point (indexed by *n*) to every line segment that connects the points along the experimental edge found for the *j*^th^ image in the time-lapse sequence, denoted *d*_*n*_. (In general, there were more points along the experimental edge than the computational edge.) The square root of the average of the squares of these minimum distances is calculated and denoted *D*_*j*_, where

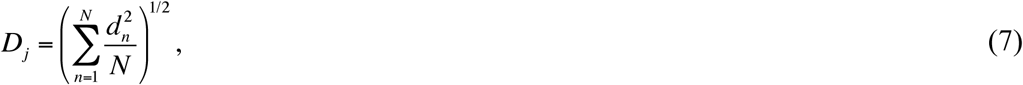

where index *n* denotes the computational points counted along the edge and *N* is the total number of these points. Summing over all time points gives the distances error term

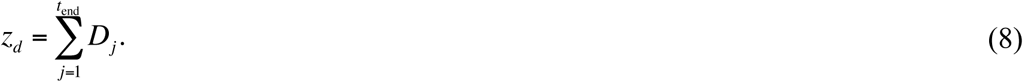

For the error in strains, we calculated the mean-squared difference between the observed and simulated density ratios. At each pixel we approximate the experimental density ratio, denoted as *ξ*_exp,*j*_, from Eq. 15 (Materials and Methods: Density Ratio Calculation via Strain Mapping Method) with the deformation gradient (Eq. 17) for the *j*^th^ (corresponding with *ρ*_*r*_) and (*j*+*δ*)^th^ (corresponding with *ρ*_*s*_) images in the time-lapse sequence, where *δ* denotes the frame increment between the pair of images. To limit the impact of inelastic mechanical events such as cell rearrangement or cell division, e.g. cases where tissue position becomes discontinuous, we limit the time between the pairs of registered images. *δ* is chosen to be the smallest frame increment in which changes between images can be detected, and in our case, *δ*=5 (S1 File). Since the size of the matrix *ξ*_exp,*j*_ may not be the same size as the computational grid (chosen for computational efficiency), we first linearly interpolate the experimental density ratio values at the computational grid nodes using the MATLAB (The MathWorks, Natick, MA) function `interp2`. The computational density ratio is denoted *ξ*_comp,*j*_, which is computed at each grid node by taking the ratio of the density from the *j*^th^ time point (corresponding with *ρ*_*r*_) and the (*j*+*δ*)^th^ time point (corresponding with *ρ*_*s*_). In other words, to compute *ξ*_comp,*j*_ where *ρ*(*x,y,t*) is the computational density for each element (*ℓ*_1_, *ℓ*_2_) in the grid,

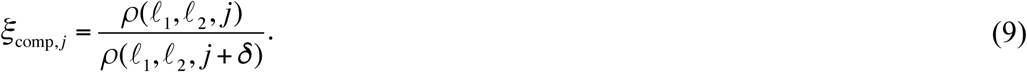

The square root of the average of the squares of the differences between the experimental and computational density ratios is

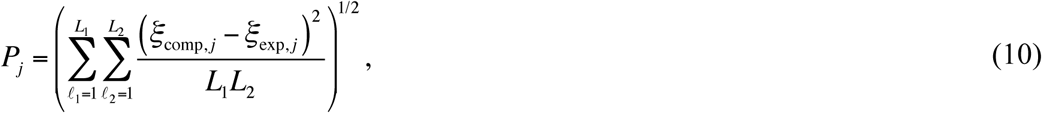

where *ξ*_comp,*j*_ and *ξ*_exp,*j*_ are evaluated at grid node (*ℓ*_1_, *ℓ*_2_), and the size of the computational grid is *L*_1_×*L*_2_. Summing over all time points (minus frame increment *δ*) gives the density ratios error term

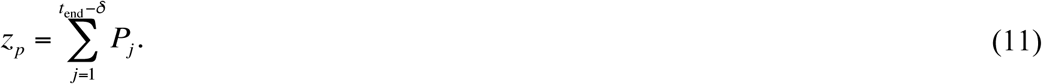

We take the total error to be

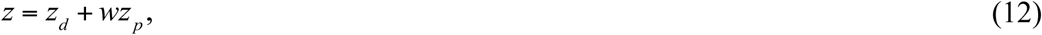

where we use weight *w* = 1000 to ensure that the distances error term *z*_*d*_ and density ratios error term *z*_*p*_ are approximately the same order. If the simulated explant migrates outside the computational domain, the error is taken to be “NaN” in our numerical code to indicate an extremely poor fit since the entire explant remains within the domain during the time-lapse sequence. If the density in the center of the explant increases, the error is also taken to be “NaN”.

### Density Ratio Calculation via Strain Mapping Method

We use edge detection to map explant boundaries and digital image correlation [43,69] of pigment patterns in *Xenopus* ectoderm to map strain rates within the spreading explant. By tracking the explant boundary and pigment patterns we produce a translation map or Eulerian description of flow can be produced showing where parcels of tissue move over time [44]. This map provides an estimate of local density change. Here, we use elastic registration to track cell movement using the pigment granule distribution in the epithelial cell sheet.

Elastic registration between two images in a time-lapse sequence separated by a fixed amount of time represents the kinematic transformation between the two images and makes minimal assumptions about the mechanical properties of the materials involved. The usefulness of the registration, i.e. that it reflects linear deformations in the ectoderm, relies on additional assumptions based on the biology of *Xenopus* embryos. Since cell division, rearrangement, and radial intercalation events are typically slow, these processes do not disrupt the pigment patterns observed at the apical face of the ectoderm.

Since we are examining a short-term analysis of strain, we assume that the *Xenopus* tissue is in a quasi-static equilibrium and mass is added very slowly between each pair of images, and thus we may assume that there is local conservation of mass. Letting *dV*_*r*_ be the infinitesimal volume in the reference material coordinates and *dV*_*s*_ be the corresponding infinitesimal volume in the spatial coordinates, the change of volume due to deformation [70] can be written as

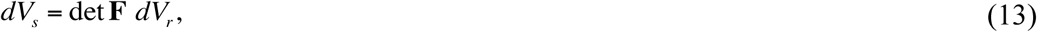

where **F** is the deformation gradient defined as 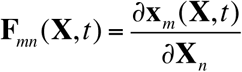, where **X**=(**X**_1_,**X**_2_) is the (*x,y*)-coordinate positions in the first still image in pixels and **x**=(**x**_1_,**x**_2_) is the (*x,y*)-coordinate positions in the second still image.

Eq. 13 implies that the Jacobian det **F** measures the ratio of the volumes. The conservation of mass equation

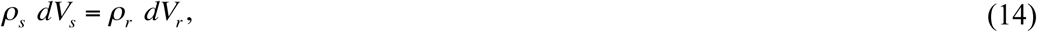

thus implies that, locally,

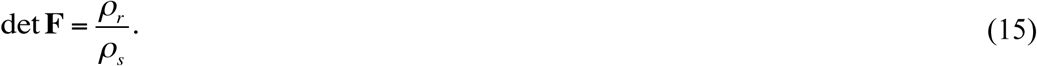

Hence, if we can calculate the deformation gradient **F**, we can obtain an estimate of the local change in density.

To perform digital image correlation on image pairs of pigmented ectoderm tissue and calculate strain, or local deformation, we use a texture mapping strategy with the ImageJ software plugin bUnwarpJ, which is used for elastic and consistent image registration ([71]; available for download at http://fiji.sc/BUnwarpJ). By using differential image correlation techniques [72], we calculate deformation of the tissue via the translation map (Fig 1E; third-fifth rows). For details on the strain mapping method discretization to obtain numerical estimates of the *x*-strain *ε*_*xx*_, *y*-strain *ε*_*yy*_, *xy*-strain *ε*_*xy*_, *yx*-strain *ε*_*yx*_ (which is equal to *ε*_*xy*_), and displacement gradient ∇**u**, please see the S1 File.

Since the gradient of the displacement vector is defined as

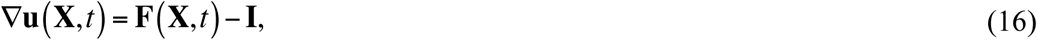

where **I** is the identity matrix, using the discretization of the displacement gradient ∇**u**(*i,j*) we numerically approximate the deformation gradient at each pixel (*i,j*) as

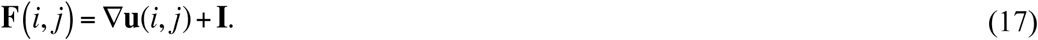

Taking the determinant for each *i* and *j*, Eq. 17 gives a numerical approximation of the density ratio in Eq. 15 at each pixel (Fig 1E; last row).

## Acknowledgements

The authors thank Deepthi Vijayraghavan for experimental contributions and David Swigon, Matthias Morzfeld, and members of the Davidson Lab for helpful discussions.

An allocation of computer time from the UA Research Computing High Performance Computing (HPC) and High Throughput Computing (HTC) at the University of Arizona is gratefully acknowledged.

## Supporting Information

**S1 File. Strain mapping method discretization.** Here we describe the implementation of the strain mapping method to calculate the deformation of a tissue via estimates of the *x*-strain *ε*_*xx*_, *y*-strain *ε*_*yy*_, *xy*-strain *ε*_*xy*_, *yx*-strain *ε*_*yx*_, and displacement gradient ∇**u** between two images in a time-lapse sequence.

**S2 File. Mathematical model derivation.** Here we describe the derivation of the mathematical model of single layer cell migration of Arciero et al. [11].

**S1 Table. 95% confidence intervals for the one-dimensional projections of the posterior distributions for each parameter for each explant.** 10,000 parameter sets were used per explant.

**S1 Fig.**
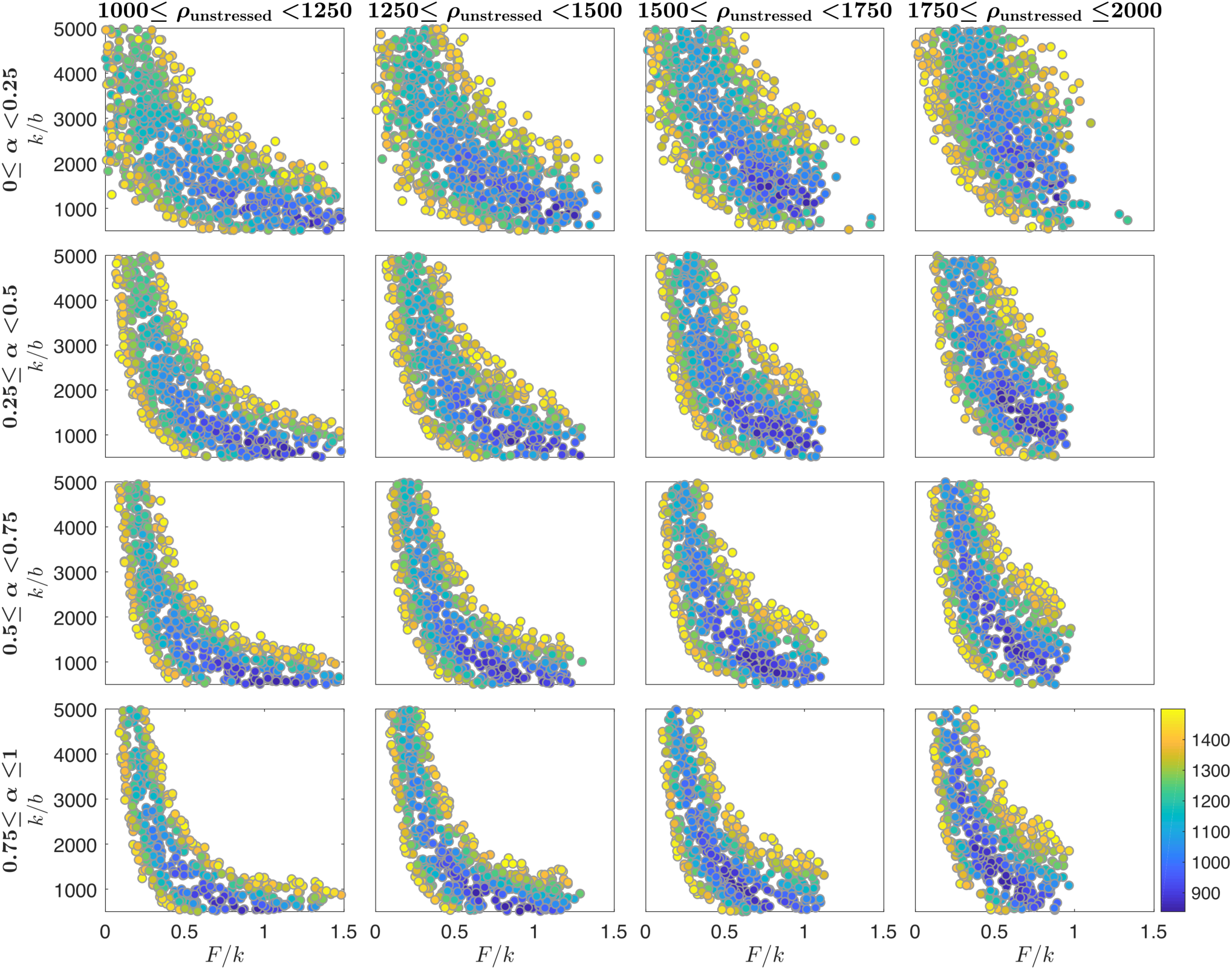
Scatterplot of accepted parameter value sets from the approximate Bayesian computation rejection method for Explant #8. Each circle indicates an accepted parameter value set found using the approximate Bayesian computation rejection method (cf. Fig 3A), and the color of the circle corresponds with its calculated error (Eq. 12).

**S2 Fig.**
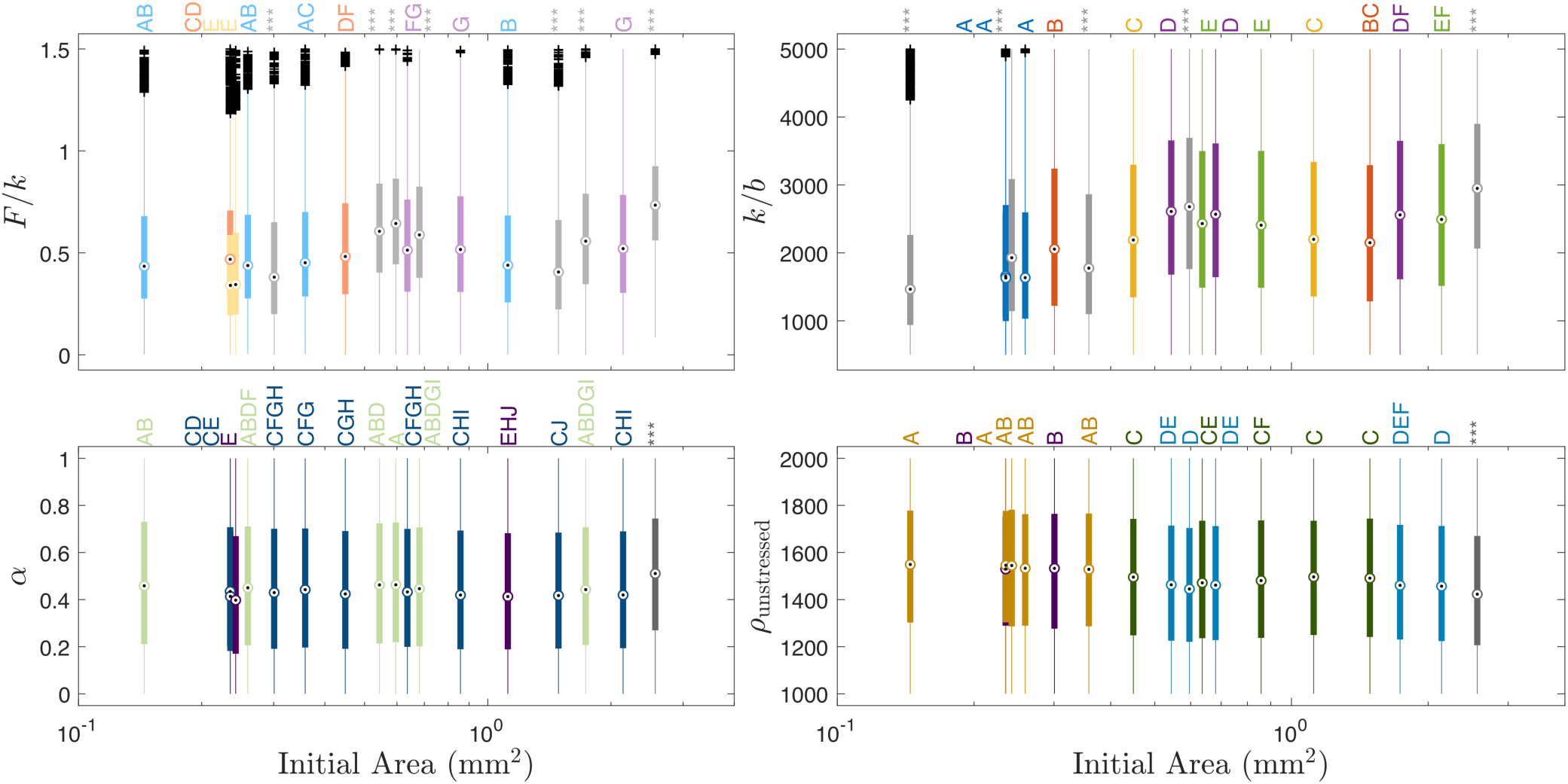
Boxplots of the 1-D projections of the posterior distributions of the parameters *F*/*k, k*/*b, α*, and *ρ*_unstressed_ for all 18 explants (cf. Fig 3C). The compact letter display (CLD) and colors reflect the results for Tukey’s multiple comparisons test; in particular, boxplots labeled with the same letter were considered not statistically different (based on significance level 0.05). Boxplots labeled with *** were statistically significantly different than all the other boxplots. Coloring was chosen to reflect groups of boxplots that were labeled similarly. The plus signs indicate outliers, the target symbols indicate the median, and the length of the boxplot indicates the interquartile range. 10,000 parameter sets were used per explant. Cf. Fig 3 for the grouped boxplots.

**S3 Fig.**
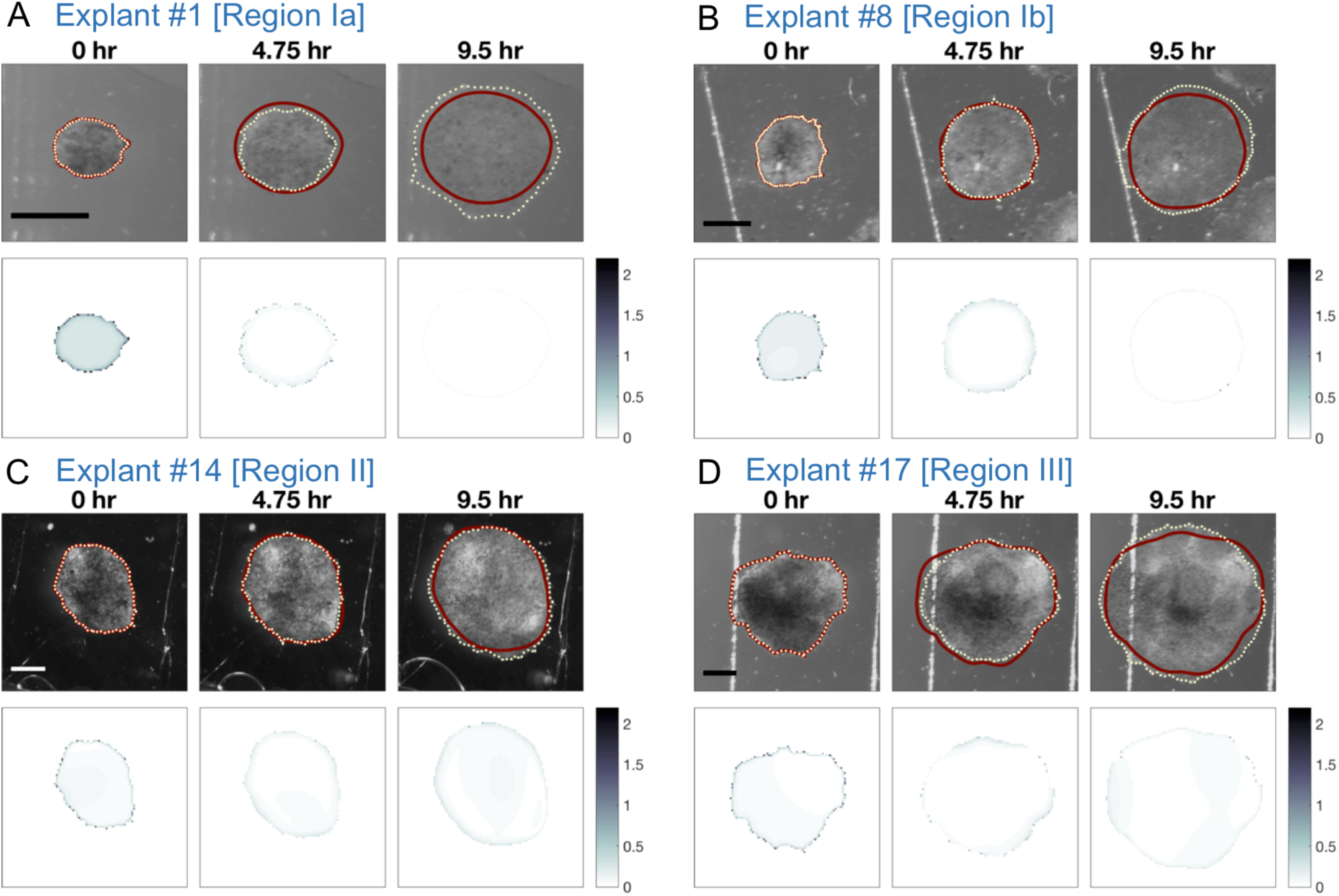
Simulation of tissue migration for Explants #1, 8, 14, and 17. Progression of *Xenopus* animal cap explant tissue migration at 95 minute time intervals for Explants #1, 8, 14, and 17 with the parameter set that resulted in the smallest error (Eq. 12). In the top panel, the computed edge from the mathematical model is represented by a solid dark red curve and the experimental edge is represented by a dotted light yellow curve. The bottom panel shows the absolute value of the density ratio at each grid node (where both density ratios are nonzero) of the experimental data between the given time point and 25 minutes later minus the computed density ratio. Scale bar: 500 µm for each explant. See the S9-12 Videos for time-lapse sequences of these still images. (A) Explant #1 [Region Ia] (initial area 0.14 mm^2^): *F/k* = 0.6101, *k/b* = 510 µm^2^/h, *α* = 0.9479 h^-1^, *ρ*_unstressed_ = 1404 cells/µm^2^. The total error with these parameters is 1122. (B) Explant #8 [Region Ib] (initial area 0.45 mm^2^): *F/k* = 0.9059, *k/b* = 510 µm^2^/h, *α* = 0.9492 h^-1^, *ρ*_unstressed_ = 1313 cells/µm^2^. The total error with these parameters is 820. (C) Explant #14 [Region II] (initial area 1.12 mm^2^): *F/k* = 0.7948, *k/b* = 635 µm^2^/h, *α* = 0.9410 h^-1^, *ρ*_unstressed_ = 1530 cells/µm^2^. The total error with these parameters is 774. (D) Explant #17 [Region III] (initial area 2.14 mm^2^): *F/k* = 0.8295, *k/b* = 950 µm^2^/h, *α* = 0.8963h^-1^, *ρ*_unstressed_ = 1752 cells/µm^2^. The total error with these parameters is 834.

**S4 Fig.**
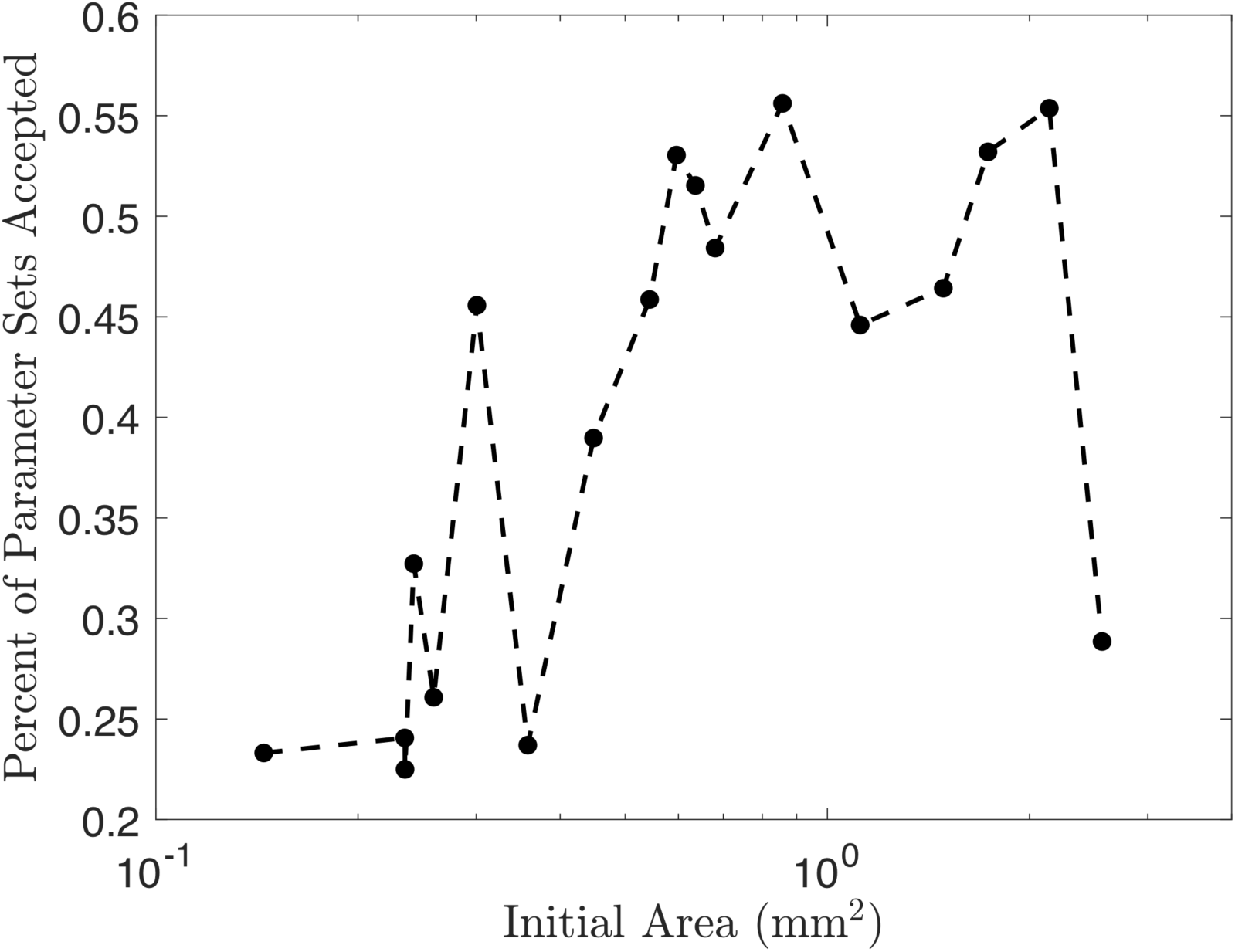
Percent of Parameter Sets Accepted in ABC Rejection Algorithm Implementation. 10,000 parameter sets that resulted in total error (Eq. 12) less than or equal to a tolerance threshold of 1500 were collected for each of the 18 explants in the model building set. More than 10,000 simulations were run to obtain 10,000 accepted parameter sets, and in general, smaller explants required more simulations run than larger explants.

**S5 Fig.**
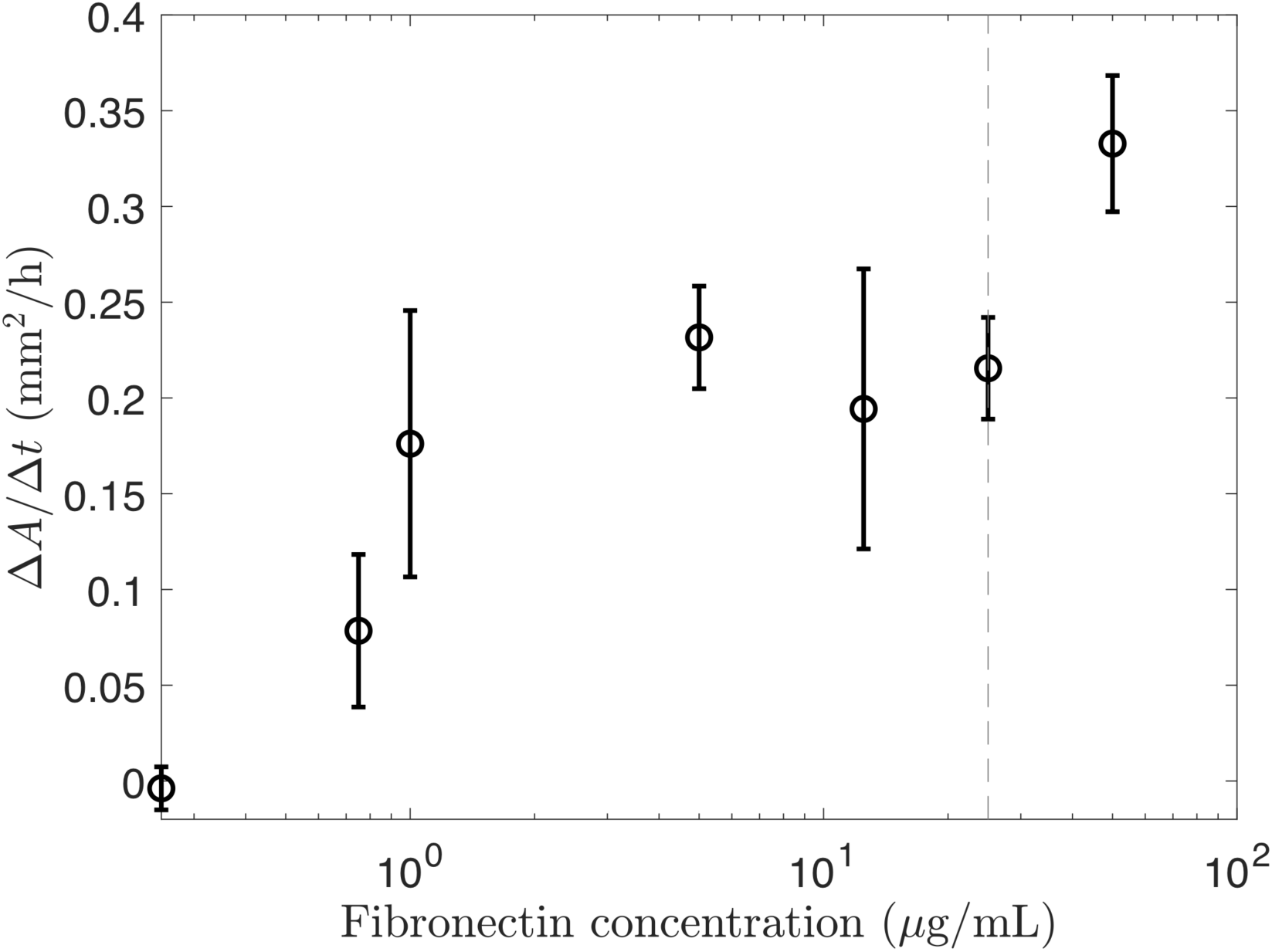
Effect of fibronectin concentration on spreading rate. Average change in area over time (Δ*A*/Δ*t*) for *Xenopus* animal cap explants that are plated on petri dishes with different concentrations of fibronectin. The error bars show the standard deviation. The spreading rate is faster for higher concentrations of fibronectin than for smaller concentrations of fibronectin. The dashed line represents the fibronectin concentration for the experiments in this paper, 25 µg/mL.

**S1 Video. Time-lapse sequences of still images in Fig 1D.**

**S2 Video. Time-lapse sequences of still images in Fig 4A.**

**S3 Video. Time-lapse sequences of still images in Fig 4B.**

**S4 Video. Time-lapse sequences of still images in Fig 4C.**

**S5 Video. Time-lapse sequences of still images in Fig 4D.**

**S6 Video. Time-lapse sequences of still images in Fig 4E.**

**S7 Video. Time-lapse sequences of still images in Fig 4F.**

**S8 Video. Time-lapse sequences of still images in Fig 4G.**

**S9 Video. Time-lapse sequences of still images in S3A Fig.**

**S10 Video. Time-lapse sequences of still images in S3B Fig.**

**S11 Video. Time-lapse sequences of still images in S3C Fig.**

**S12 Video. Time-lapse sequences of still images in S3D Fig.**

### Strain mapping method discretization

Here we describe the implementation of the strain mapping method to calculate the deformation of a tissue via estimates of the *x*-strain *ε*_*xx*_, *y*-strain *ε*_*yy*_, *xy*-strain *ε*_*xy*_, *yx*-strain *ε*_*yx*_, and displacement gradient ∇**u** between two images in a time-lapse sequence.

For images of width *n* pixels and height *m* pixels, the properties of each pixel can be represented by an entry in an *m*×*n* matrix. For each pair of images from the time-lapse sequence, let **X**=(**X**_1_,**X**_2_) be the (*x,y*)-coordinate positions in the first still image in pixels and **x**=(**x**_1_,**x**_2_) be the (*x,y*)-coordinate positions in the second still image. The top left corner of an image is the origin, and *x* increases from left to right and *y* increases from top to bottom. The entries of **X** are therefore defined by **X**_1_(*i,j*)=*j*-1 and **X**_2_(*i,j*)=*i*-1 for *i*=1,2,…,*n*, and *j*=1,2,…,*m*.

We mask the pair of images so that registration occurs only in the actual location of cells. We then initialize bUnwarpJ to calculate the coefficients of the cubic B-spline map *β* that defines the transformation 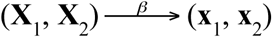 [1]. Initializing bUnwarpJ again, we apply *β* to **X** by converting the transformation to “raw” data, which reports the elastically-mapped position of each pixel in the first image to the “fitted” position in the second image. We obtain **x**, the mapped position of each pixel from the first image to its position in the second image, in pixels. Note that pixels outside of the mask will be mapped as well, but we will remove this extraneous data after we have obtained the strains. Next, we calculate the displacement vector **u** by

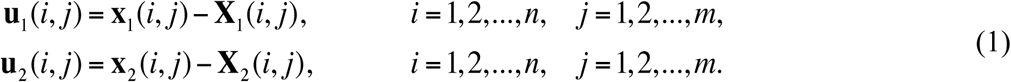

The engineering, or Cauchy, strain is defined as

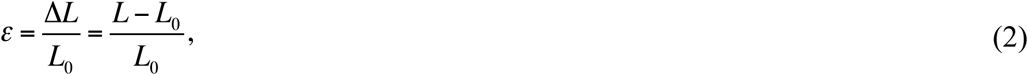

where Δ*L* is the change in length of the tissue, *L*_0_ is the original length, and *L* is the current length. The displacement vector **u** is converted into *x*-strain, *y*-strain, *xy*-strain, and *yx*-strain by

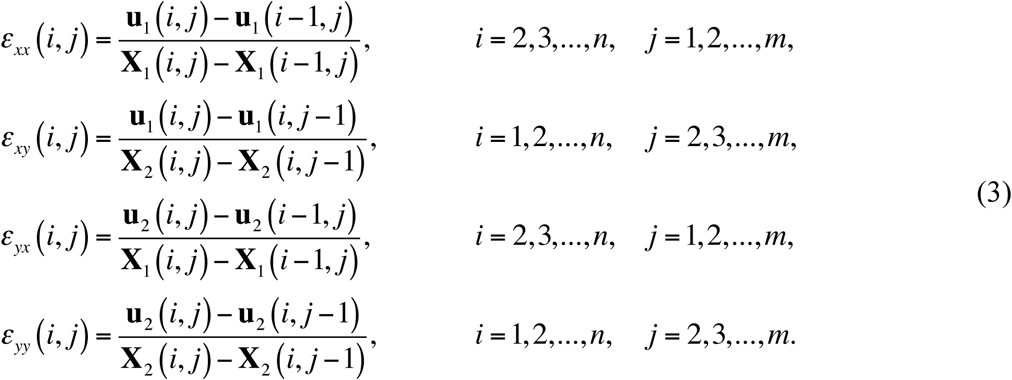

Note that all of the denominators above equal 1 pixel and the shear strains *ε*_*xy*_ = *ε*_*yx*_. At this point, the strains can be visualized to show where there are contractions in the tissue (*ε* < 0) and where there are dilations (*ε* > 0) [2].

Using these strain calculations, we can numerically approximate the displacement gradient at each pixel as

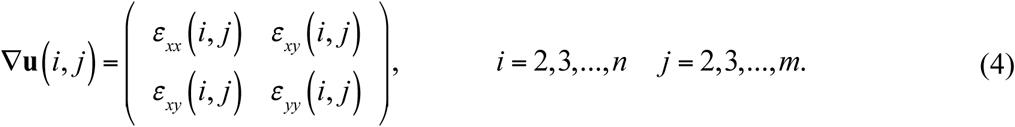

We found that we could limit numerical boundary effects on the registration by ensuring the tissue was at least 200 pixels from the outer boundary of the image. To ensure the registration between images was detecting movement and not noise, we chose a time interval long enough for movement to be discernible. In our case, a time interval of 25 minutes ensured that the relative change in area between image pairs was on average more than 5%, which would correspond with strain measurements above the noise between images.

## Mathematical model derivation

Here we describe the derivation of the mathematical model of single layer cell migration of Arciero et al. [1].

The cell layer is represented as a 2D compressible fluid, and the variable *ρ* describes the tissue density as a function of position **x**=(*x,y*) and *t*. The law of conservation of mass,

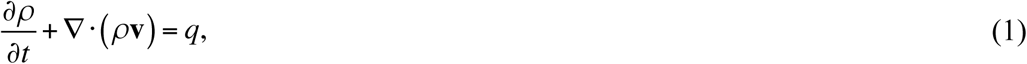

where **v** is the velocity of the cell layer, includes the growth term *q* which may generally depend on space **x**, time *t*, or density *ρ*, and describes the net rate of change in the number of cells within the layer.

Balance of linear momentum implies

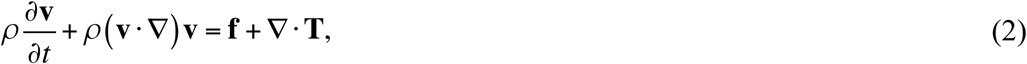

where the tensor **T** represents the stresses within the cell layer and **f** accounts for the force of adhesion of the cell layer to the substrate. **f** is the result of the action exerted on a material element by the substrate, i.e. the negative of traction force. It is assumed that the **f** is negatively proportional to the cell layer velocity,

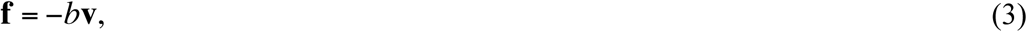

where b is a constant of adhesion. The cell layer is assumed to behave as a compressible inviscid fluid with the constitutive equation

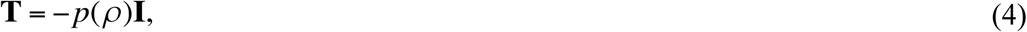

where *p* is the pressure within the cell layer. The pressure depends on the tissue density and is taken to be positive when cells are compressed and negative when cells are stretched. Assuming acceleration is negligible and substituting Eqs. 3 and 4 into Eq. 2 we obtain the equation

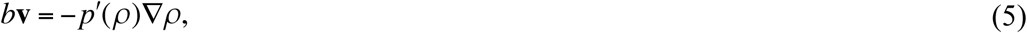

which is the relation between the velocity of cells and the gradient of tissue density; it resembles Darcy’s law describing the flow of fluid through a porous medium.

Substituting Eq. 5 into Eq. 1 results in the governing equation that describes the evolution of tissue density,

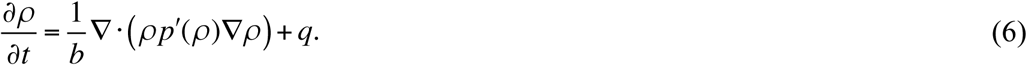

Given the presence of lamellipodia on the edge of the ectoderm, i.e. tissue boundary 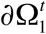, we assume there is a constant force per unit length *F* (see Table 1 and Fig 2A-B) exerted outward at the tissue boundary that is equal in magnitude to that of the force of the cells in the interior. To express this boundary condition in mathematical notation, a function describing the forces within the tissue is necessary. In Arciero et al. [1], various constitutive relations for function *p*(*ρ*) were considered, but the main relation studied was

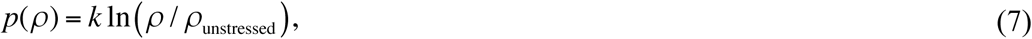

where *ρ*_unstressed_ is a parameter described in Table 1, as it gave appropriate behavior at both large and small densities. Substituting the constitutive relation in Eq. 7 into Eq. 6 gives the governing equation

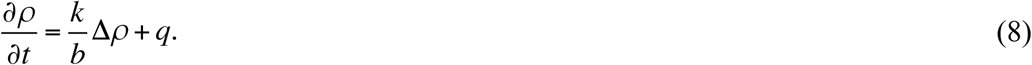

Notice that the Laplacian that appears in the governing equation is due to the constitutive relation chosen for the pressure and hence the governing equation should not be thought of as reaction-diffusion equation. The Laplacian does not arise from any underlying diffusion process or Brownian motion and *k/b* should not be interpreted as a diffusion constant (Arciero et al. [1]).

One boundary condition imposed on the boundary of the cell layer is that there is a constant force per unit length *F* outward directed against the substrate due to lamellipodia, which requires setting *p* = -*F* at the boundary [1,2] in Eq. 7 and then solving for *ρ*, resulting in

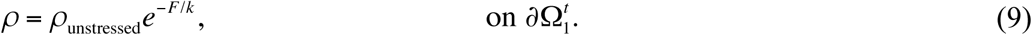

Another boundary condition imposed on the boundary of the cell layer is a Stefan condition, which describes the speed of the moving edge. This condition comes from Eq. 5 evaluated at Eq. 9 and is

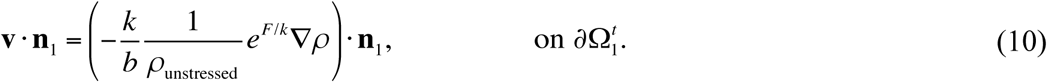

where **v**(**x**,*t*) is the velocity of the layer and **n**_1_(**x**,*t*) is the outward unit normal to the tissue boundary ∂Ω_1_^t^

On the edge of the computational domain ∂Ω_2_, we assume that there is no flux of cells, i.e. cells are unable to move beyond this boundary, and so we have the Neumann boundary condition

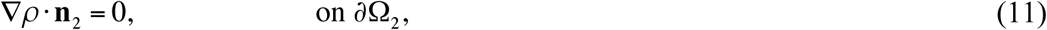

where **n**_2_(**x**,*t*) is the outward unit normal to the edge of the computational domain ∂Ω_2_.

By segmenting cells in confocal images of the epithelial layer of a representative animal cap explant labeled with a GFP-membrane tag, we measured the tissue density in the epithelial layer at the initial imaging time point to be 0.0047 cells/µm^2^, so we take the initial condition to be

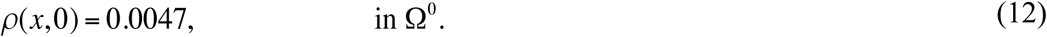

---

| Parameter | | $F/k$ (dimensionless) | | $k/b$ ($\mu\text{m}^2/\text{h}$) | | $\alpha$ ($\text{h}^{-1}$) | | $\rho_{\text{unstressed}}$ (cells/$\mu\text{m}^2$) | |
| --- | --- | --- | --- | --- | --- | --- | --- | --- | --- |
| Explant | Initial Area ($\text{mm}^2$) | Lower Bound | Upper Bound | Lower Bound | Upper Bound | Lower Bound | Upper Bound | Lower Bound | Upper Bound |
| 1 | 0.14 | 0.4949 | 0.5060 | 1700 | 1739 | 0.4678 | 0.4793 | 1528 | 1539 |
| 2 | 0.23 | 0.5188 | 0.5299 | 1904 | 1946 | 0.4514 | 0.4629 | 1513 | 1524 |
| 3 | 0.23 | 0.4112 | 0.4223 | 1937 | 1983 | 0.4376 | 0.4491 | 1528 | 1539 |
| 4 | 0.24 | 0.4156 | 0.4269 | 2161 | 2209 | 0.4244 | 0.4358 | 1523 | 1535 |
| 5 | 0.26 | 0.4980 | 0.5091 | 1903 | 1946 | 0.4585 | 0.4699 | 1519 | 1530 |
| 6 | 0.30 | 0.4397 | 0.4517 | 2262 | 2311 | 0.4470 | 0.4584 | 1512 | 1524 |
| 7 | 0.36 | 0.5084 | 0.5195 | 2047 | 2093 | 0.4504 | 0.4618 | 1515 | 1526 |
| 8 | 0.45 | 0.5309 | 0.5425 | 2355 | 2402 | 0.4419 | 0.4532 | 1491 | 1502 |
| 9 | 0.54 | 0.6276 | 0.6391 | 2653 | 2700 | 0.4658 | 0.4772 | 1468 | 1479 |
| 10 | 0.60 | 0.6630 | 0.6741 | 2718 | 2764 | 0.4697 | 0.4811 | 1460 | 1471 |
| 11 | 0.64 | 0.5430 | 0.5549 | 2508 | 2556 | 0.4497 | 0.4610 | 1480 | 1491 |
| 12 | 0.68 | 0.6079 | 0.6197 | 2624 | 2672 | 0.4545 | 0.4659 | 1467 | 1478 |
| 13 | 0.86 | 0.5499 | 0.5621 | 2502 | 2550 | 0.4423 | 0.4538 | 1482 | 1494 |
| 14 | 1.12 | 0.4848 | 0.4965 | 2366 | 2415 | 0.4357 | 0.4470 | 1489 | 1500 |
| 15 | 1.49 | 0.4585 | 0.4704 | 2318 | 2366 | 0.4397 | 0.4510 | 1488 | 1499 |
| 16 | 1.73 | 0.5779 | 0.5898 | 2614 | 2663 | 0.4550 | 0.4663 | 1469 | 1481 |
| 17 | 2.14 | 0.5524 | 0.5649 | 2556 | 2605 | 0.4419 | 0.4532 | 1464 | 1476 |
| 18 | 2.57 | 0.7522 | 0.7621 | 2943 | 2987 | 0.5011 | 0.5122 | 1440 | 1451 |

## Notes

#### Summary of Updates

Copyediting updates.

https://dx.doi.org/10.5061/dryad.8pj52vk

https://github.com/tstepien/strain-mapping

https://github.com/tstepien/cell-migration-spatialcoord-onelayer

